# Ancestral heterogeneity of ancient Eurasians

**DOI:** 10.1101/268524

**Authors:** Daniel Shriner

## Abstract

Supervised clustering or projection analysis is a staple technique in population genetic analysis. The utility of this technique depends critically on the reference panel. The most commonly used reference panel in the analysis of ancient DNA to date is based on the Human Origins array. We previously described a larger reference panel that captures more ancestries on the global level. Here, we reanalyzed DNA data from 279 ancient Eurasians using our reference panel. We found substantially more ancestral heterogeneity than has been reported. Our reanalysis provides evidence against a resurgence of Western hunter-gatherer ancestry in the Middle to Late Neolithic and evidence for a common ancestor of farmers characterized by Western Asian ancestry, a transition of the spread of agriculture from demic to cultural diffusion, at least two migrations between the Pontic-Caspian steppes and Bronze Age Europe, and a sub-Saharan African component in Natufians that localizes to present-day southern Ethiopia.

---

Before the technological advances that permitted ancient DNA studies, historical inferences were made using present-day samples in conjunction with well-established theory and techniques in population genetics and phylogenetic reconstruction. In particular, inferences regarding population structure have been based on the popular software STRUCTURE^1^, ADMIXTURE^2^, and variants thereof. The basic idea is to perform model-based estimation of ancestries from multi-locus genotypes.

Having learned ancestry-specific allele frequencies in unsupervised clustering analysis from a data set, it is computationally efficient to project new samples onto the ancestries in order to learn about population structure in the new samples. The utility and quality of projection analysis, or supervised clustering analysis, strongly depends on the reference set of learned ancestries. In the analysis of ancient DNA, the reference panel most widely used to date comprises <3,000 individuals^3–6^, although some data are not freely available. One consequence of widespread use of this single reference panel is consistency within the ancient DNA field. Unfortunately, the labels for ancestries used in these papers lacks overlap with labels used by other researchers for ancestries in present-day peoples. Furthermore, none of the results in the ancient DNA papers has been replicated using a second reference panel. Here, we combined completely public domain data to generate a reference panel comprising 5,966 individuals from 282 samples, from which we estimated 21 ancestries^7^. After projecting 279 ancient Eurasians onto our reference panel, we reached a distinct series of conclusions regarding the genetic history of Europe and Western Asia.

## Results

### European Hunter-Gatherers

First, we considered the Western (Hungary, Luxembourg, Spain, and Switzerland), Scandinavian (Sweden), and Eastern (Karelia and Samara) hunter-gatherers (Figure 1A). These 14 hunter-gatherers averaged 71.6% Northern European ancestry, 27.4% Southern European ancestry, and 0.9% Oceanian ancestry (Table 1). The Y DNA haplogroups included eight I2, one C1a2, one J, one R1a, and one R1b (Supplementary Table S1). The mitochondrial haplogroups included 13 U (including nine U5), one C, and one R (Supplementary Table S1). In contrast to previous reports, Western and Eastern hunter-gatherers were not homogeneous for different ancestries^8^ nor were they separated^9^. Rather, Amerindian, Circumpolar, and Southern Asian ancestries decreased from east to west, complemented by an increase of Southern European ancestry from east to west (Table 1). Thus, these three ancestries existed in Eastern and, to a lesser extent, Scandinavian hunter-gatherers thousands of years before the European Bronze Age and in higher proportions than in the Bronze Age steppe populations.

**Table 1.**
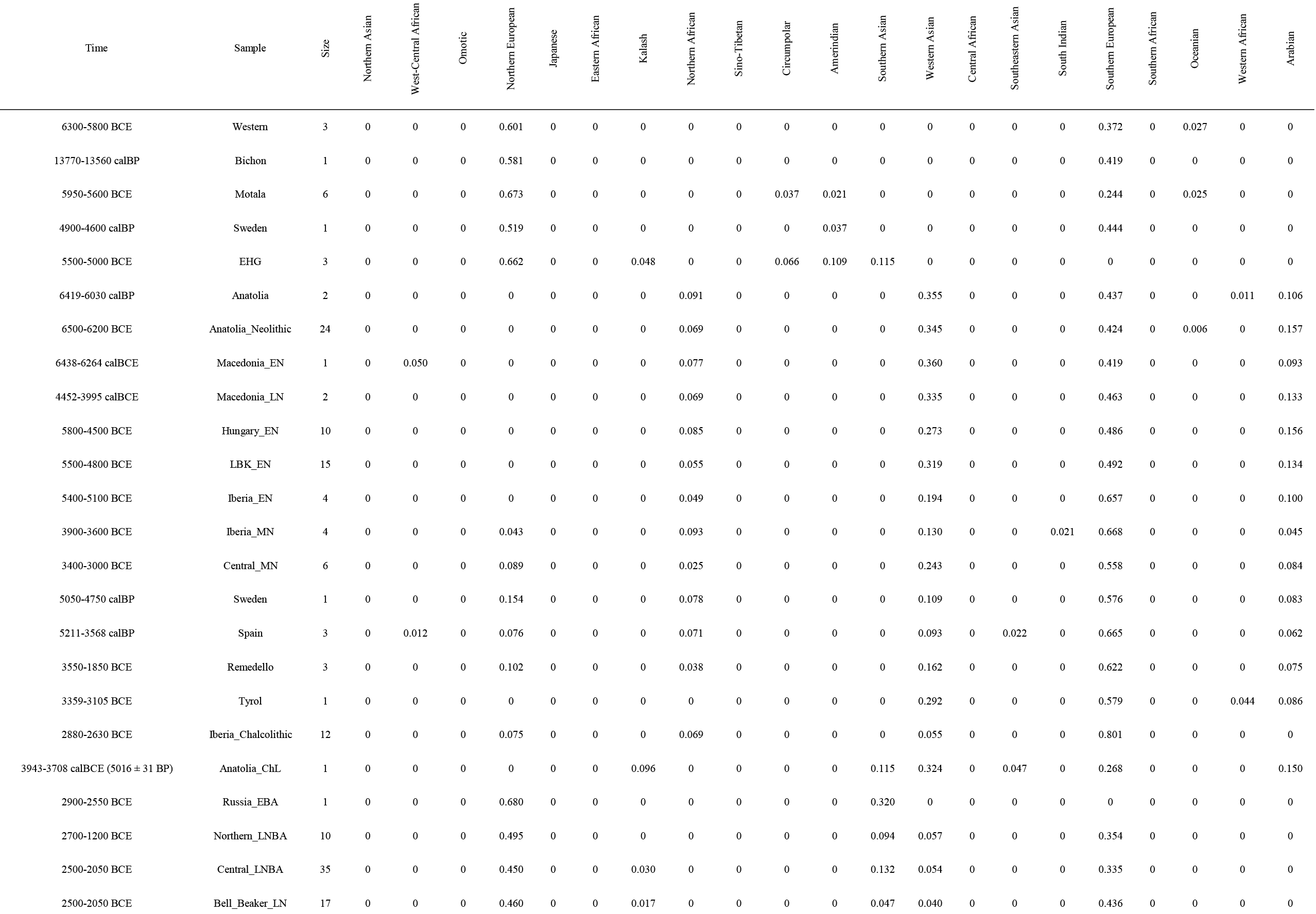
Mean ancestry proportions for 49 samples of ancient Eurasians.

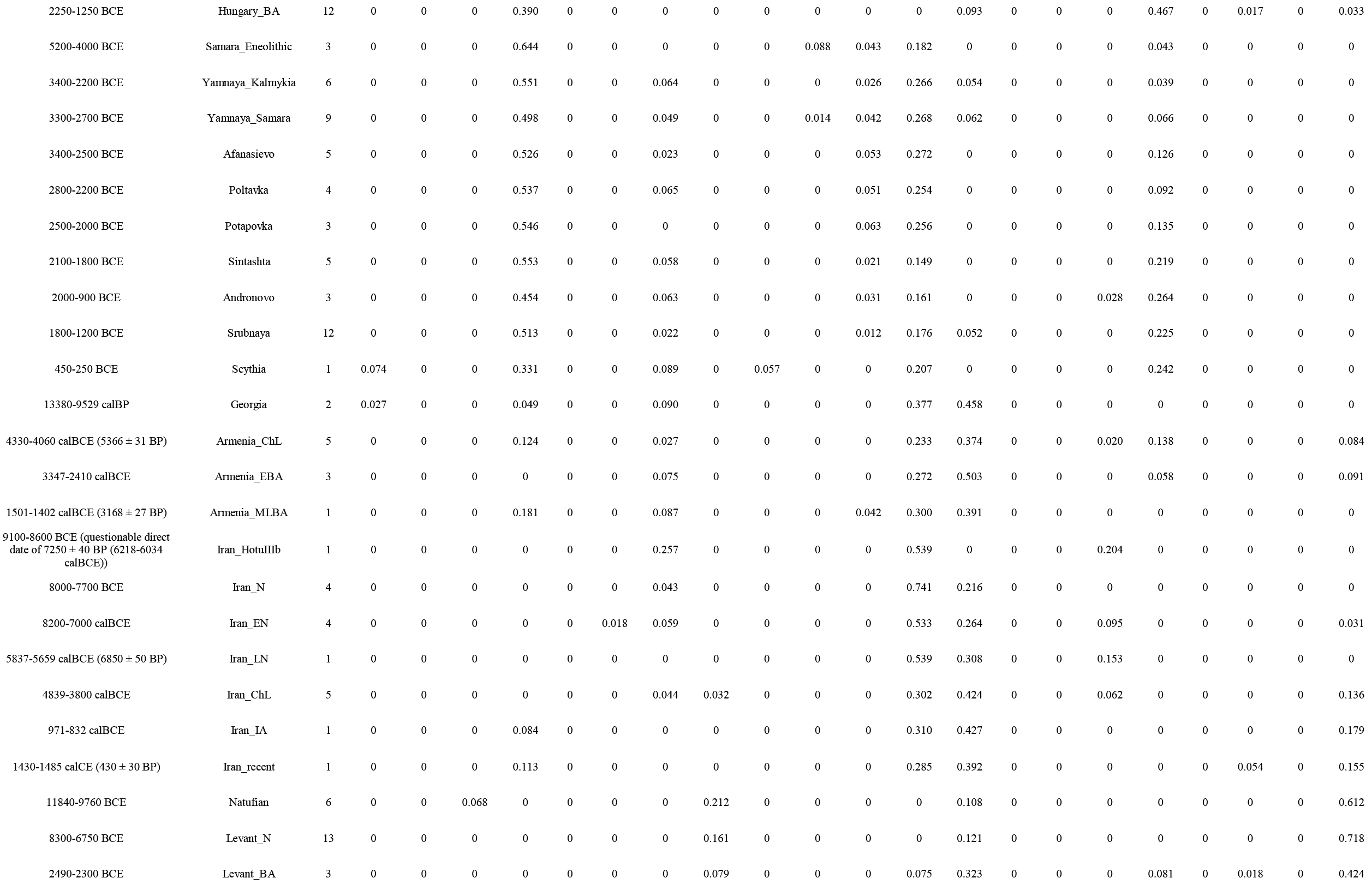

**Figure 1.**
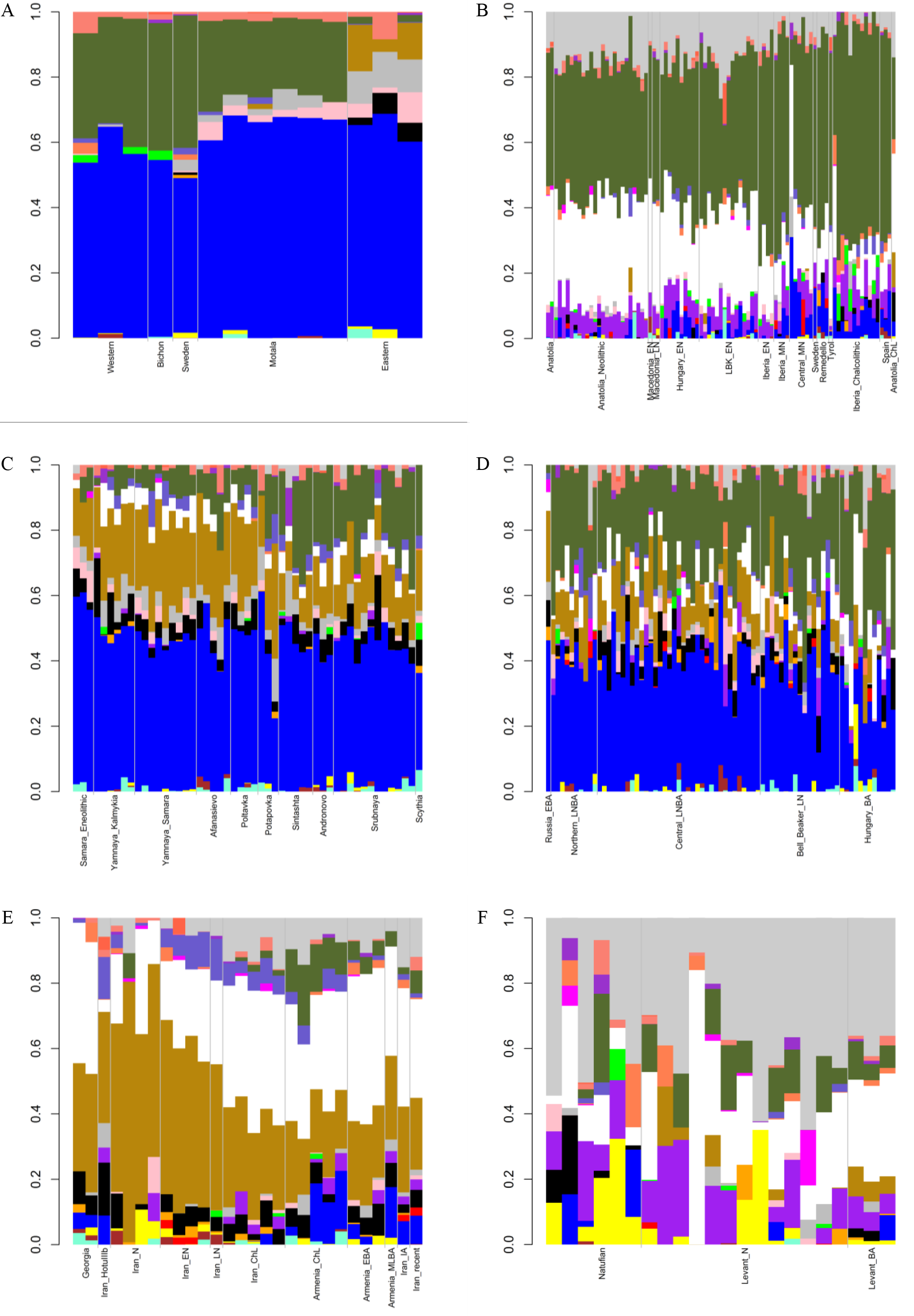
Admixture bar plots showing projections of ancient Eurasians onto 21 ancestries. The proportions are the raw output from ADMIXTURE. The 21 ancestral components are Southern African (dark orchid), Central African (magenta), West-Central African (brown), Eastern African (orange), Omotic (yellow), Northern African (purple), South Indian (slate blue), Kalash (black), Japanese (red), Sino-Tibetan (green), Southeastern Asian (coral), Northern Asian (aquamarine), Amerindian (gray), Oceanian (salmon), Southern European (dark olive green), Northern European (blue), Western Asian (white), Arabian (light gray), Western African (tomato), Circumpolar (pink), and Southern Asian (dark goldenrod). (A) Hunter-gatherers. (B) Early Neolithic to Copper Age Europeans. (C) Steppe peoples. (D) Bronze Age Europeans. (E) Western Asians. (F) Levantines.

The two Georgian hunter-gatherers did not group with the European hunter-gatherers. The Georgian hunter-gatherers were predominantly a mixture of 45.8% Western Asian and 37.7% Southern Asian ancestries, with only 4.9% Northern European and no Southern European ancestries (Table 1). The Y DNA haplogroups were J and J2 (Supplementary Table S1). These results support an association between Y DNA haplogroup J2 and either Western Asian ancestry^10,11^ or Southern Asian ancestry. The mitochondrial haplogroups H and K in the Georgian hunter-gatherers were not observed in the European hunter-gatherers (Supplementary Table S1).

### Early Neolithic Peoples

Second, we considered the Early Neolithic samples from Germany, Hungary, and Iberia, collectively referred to early European farmers, as well as from Anatolia and Macedonia (Figure 1B). These 56 individuals averaged 47.2% Southern European, 31.9% Western Asian, 14.2% Arabian, and 6.8% Northern African ancestries (Table 1). The Y DNA haplogroups included 13 G, four H, four I, three C, two T, one J, and one R1b (Supplementary Table S1). Western Asian ancestry currently co-localizes in the Caucasus with Y DNA haplogroup G; the 46.4% frequency of the G haplogroup was not different from the autosomal proportion of Western Asian ancestry (*p* = 0.148). Multiple descendants of the mitochondrial haplogroup R (H, V, J, T, U, and K) accounted for 69.6% of lineages (Supplementary Table S1). Thus, the Early Neolithic samples qualitatively differed from hunter-gatherers by harboring more diverse sets of Y DNA haplogroups and mitochondrial lineages.

### Early to Middle Bronze Age Steppe Peoples

Third, we considered the Eurasian steppe peoples (Figure 1C). The Eneolithic Samara sample had 64.4% Northern European, 18.2% Southern Asian, 8.8% Circumpolar, 4.3% Amerindian, and 4.3% Southern European ancestries (Table 1). The 27 Early to Middle Bronze Age steppe individuals (Yamnaya from Kalmykia, Yamnaya from Samara, Afanasievo, Poltavka, and Potapovka) averaged 54.7% Northern European, 27.8% Southern Asian, 7.9% Southern European, 4.7% Kalash, 4.2% Amerindian, and 0.8% Western Asian ancestries (Table 1). We included the Potapovka sample here because the sum of absolute differences in ancestry was greater post-Potapovka rather than post-Poltavka. The increases in Southern Asian and Southern European ancestries do not fit with a European hunter-gatherer source^5^ and more broadly do not fit with any of the samples, suggesting an unknown source population. Currently, Southern Asian ancestry co-localizes with Y DNA haplogroup L and correlates with Indo-Iranian languages^7^.

Although there are no L haplogroups in any of these Early to Middle Bronze Age steppe individuals (Supplementary Table S1), the correlation with Indo-Iranian languages strengthens the connection between Early to Middle Bronze Age steppe peoples and the introduction of Indo-European languages into Europe. In the Early to Middle Bronze Age steppe peoples, 83.3% of Y DNA haplogroups were R1b and 85.2% of mitochondrial haplogroups were H, J, T, or U (Supplementary Table S1). Thus, Northern European ancestry was primarily associated with R1b in these peoples, rather than with I2 as in the European hunter-gatherers, while the mitochondrial lineages were more diverse than in the European hunter-gatherers but less diverse than in the Early Neolithic peoples.

### Middle to Late Neolithic European Peoples

Fourth, we considered the Middle to Late Neolithic European peoples (Figures 1B and 1D). The 10 Middle Neolithic individuals averaged 64.2% Southern European, 18.2% Western Asian, 6.2% Northern European, 6.2% Arabian, 4.3% Northern African, and 0.9% Oceanian ancestries (Table 1). Five of six Y DNA haplogroups were G, H, or I; the one other haplogroup was R (Supplementary Table S1). The mitochondrial haplogroups were H, HV, K, T, or U (Supplementary Table S1). The 21 Copper Age individuals averaged 71.8% Southern European, 10.9% Western Asian, 7.6% Northern European, 5.6% Northern African, and 4.2% Arabian ancestries (Table 1). The Y DNA haplogroups were 75.0% I (Supplementary Table S1). This time period includes the Tyrolean Iceman who had 57.9% Southern European and 29.2% Western Asian ancestries (Table 1), the latter of which is consistent with his Y DNA haplogroup G2a^12^. Thus, as late as the Copper Age, Southern European ancestry was associated predominantly with haplogroup I. The mitochondrial haplogroups were 85.7% H, U, K, or J (Supplementary Table S1). The transition from Middle Neolithic to Copper Age involved the acquisition of 7.6% Southern European, 1.4% Northern European, and 1.3% Northern African ancestries. While Northern African ancestry increased, Arabian ancestry decreased, possibly indicating entry into Europe from northwest Africa rather than northeast Africa. The ratio of 5.4-fold more Southern European than Northern European ancestries and the presence of Northern African ancestry acquired from the Early Neolithic to the Copper Age are inconsistent with a resurgence of peoples related to Western hunter-gatherers, given that Western hunter-gatherers had 1.6-fold more Northern European than Southern European ancestry and no Northern African ancestry. Instead, this ancestral profile is suggestive of an expansion of peoples from Southern Europe resembling those from the Remedello culture. The 75 Bronze Age individuals averaged 49.0% Northern European, 40.0% Southern European, 5.9% Western Asian, and 5.1% Southern Asian ancestries (Table 1). The transition from the Copper Age to the Bronze Age involved the acquisition of 41.4% Northern European and 5.1% Southern Asian ancestries. A low level of Y DNA haplogroup R had been present in Europe for thousands of years; in the Bronze Age, however, the Y DNA haplogroups increased to 48.0% R (10 R1a and 10 R1b), while the mitochondrial haplogroups remained 90.6% H, V, K, U, J, or T (Supplementary Table S1), consistent with male-biased gene flow^13^.

### Late Bronze Age Steppe Peoples

The 20 Late Bronze Age steppe individuals (from the Andronovo, Sintashta, and Srubnaya samples) had 51.5% Northern European, 23.5% Southern European, 17.1% Southern Asian, 3.2% Western Asian, 3.1% Kalash, and 1.6% Amerindian ancestries (Figure 1C and Table 1). Thus, post-Potapovka, population change in the steppes involved an increase of 15.6% Southern European and 2.4% Western Asian ancestries (a ratio of 6.5) and a decrease of 10.7% Southern Asian, 3.2% Northern European, and 2.6% Amerindian ancestries. This change does not fit with gene flow of people like the Early Neolithic peoples^5^, who had a ratio of Southern European to Western Asian ancestry of 1.5; the source is more consistent with European Copper or Bronze Age peoples, in whom the ratios were 6.6 and 6.8, respectively. In contrast to the Early to Middle Bronze Age steppe samples, all eight Y DNA haplogroups were R1a, while 90% of mitochondrial haplogroups were H, V, U, K, J, or T (Supplementary Table S1). Collectively, these findings suggest two distinct male-biased migrations from the steppes to Europe, rather than one prolonged event^13^. The first event was associated with Early to Middle Bronze Age steppe peoples located north of the Black Sea and characterized by R1b; this incoming ancestry is associated with present-day Southern European ancestry. The second event was associated with Late Bronze Age steppe peoples located north and east of the Caspian Sea and characterized by R1a; this incoming ancestry is associated with present-day Northern European ancestry. There was also gene flow from Europe to the steppes associated with the transition from the Middle Bronze Age to the Late Bronze Age.

The Amerindian and Circumpolar ancestries were shared with the Eastern hunter-gathers from Karelia and Samara as well as all the steppe peoples but were absent from the other Europeans from the Early Neolithic through the Bronze Age. The proportion of Amerindian ancestry decreased with time, suggesting a shared relationship before the Neolithic. The ancestry shared between the steppe populations and the Caucasian hunter-gatherers was predominantly Southern Asian, not Western Asian.

### Western Asian Peoples

The ancient Iranians were characterized through the Late Neolithic period by predominantly Southern Asian ancestry (Figure 1E and Table 1). The proportion of Western Asian ancestry doubled through the Iron Age, suggesting gene flow from the Caucasus rather than the Levant^8^, while smaller amounts of Arabian and South Indian ancestries suggest gene flow from the west and the east, respectively. The ancient Armenians resembled the Georgian hunter-gatherers in having a mixture of Western Asian and Southern Asian ancestries (Figure 1E and Table 1). Northern European, Southern European, and Arabian ancestries increased throughout the Copper and Bronze Ages, again suggesting gene flow from multiple directions.

The Natufian sample consisted of 61.2% Arabian, 21.2% Northern African, 10.9% Western Asian, and 6.8% Omotic ancestry (Figure 1F and Table 1). Previously, no significant sharing of ancestral components with sub-Saharan African populations was found to accompany the presence of E1b1b1b2 Y haplogroups^8^. E1b1b1b1a-M81, but not E1b1b1b2-Z830, is presently common among Berbers in North Africa^14^. However, E1b1b1b1a-M81 has a time to most recent common ancestor of only 2,300 (95% confidence interval [1900,2700]) years before present^15^ and therefore was not prevalent in Northern African ancestry during the Epipaleolithic. Ancestry shared by Omotic-speaking peoples is found predominantly in present-day southern Ethiopia and is associated with haplogroup E, thus revealing a plausible source. The transition in the Levant from the Epipaleolithic to the Neolithic period involved an increase of Arabian ancestry at the expense of Northern African and Omotic ancestries. The transition from the Neolithic period to the Bronze Age involved the acquisition of principally Western Asian ancestry, with smaller contributions of Southern European and Southern Asian ancestries. Lazaridis *et al*.^8^ suggested that this change resulted from admixture from people resembling Chalcolithic Iranians. This putative source is unlikely because none of the ancient Iranian samples had Southern European ancestry; a Caucasian source, such as the Chalcolithic or Early Bronze Age Armenians, provides a better fit.

## Discussion

Using a large, global reference panel, we found more population structure than previously reported among 279 ancient Eurasians. All samples showed multiple autosomal ancestries, Y DNA haplogroups, and mitochondrial haplogroups. Given such a large amount of ancestral heterogeneity, previous estimates of allele frequencies, including claims of natural selection, may have been confounded by this unrecognized population structure.

The early European farmers had no Southern Asian ancestry, which does not support an origin in the eastern part of Western Asia, *i.e.*, present-day Iran. However, ancient Western Asian peoples and early European farmers shared Western Asian ancestry, and thus were not genetically dissimilar^9^. The increase of Western Asian ancestry in the Bronze Age Levant and throughout Neolithic Western Asia is consistent with demic diffusion of agriculture via a single origin, with the original people characterized by Western Asian ancestry. Even if farming was introduced into Europe by such individuals, then subsequent migrations of semi-nomadic pastoralists from the steppes during Bronze Age Europe suggests that the ultimate spread of agriculture occurred by cultural, not demic, diffusion.

We found evidence of autosomal ancestries implicating populations ranging from Central Asia, North Asia, South Asia, Northern Africa, the Middle East, and the Caucasus in contributing to the genetic origins of Bronze Age Europeans. Ancient Eurasians from the steppes had Southern Asian ancestry with little to no Western Asian ancestry. Furthermore, the shared ancestry between the Caucasian hunter-gatherers and the steppe peoples was predominantly Southern Asian. At present, Southern Asian ancestry correlates with the Indo-Iranian branch of the Indo-European language family whereas Western Asian ancestry correlates with the Ibero-Caucasian language family^7^. Taken together, it is unlikely that the Caucasian hunter-gatherers represent a primary source population of the steppe peoples, as previously suggested^10^.

Using TreeMix^16^ to reconstruct migration graphs from ancestries inferred by ADMIXTURE, we previously observed that Southern European and Northern European ancestries clustered with 77% probability and that Southern European and Arabian ancestries clustered with 23% probability^17^. We hypothesized that the primary mode reflected the relationship between R1a, characteristic of present-day Northern European ancestry, and R1b, characteristic of present-day Southern European ancestry. We further hypothesized that the secondary mode reflected the relationship between I2, present in lower frequencies in present-day Southern European ancestry, and J (more precisely, J1), characteristic of Arabian ancestry. Our current findings support both hypotheses. The fact that Southern European ancestry experienced a replacement of haplogroup I by haplogroup R and yet was inferred by ADMIXTURE to be one ancestry, rather than two distinct ancestries, serves as a strong caveat in the interpretation of ancestries, while TreeMix could detect both stages of Southern European ancestry.

All the ancestries in our reference panel were estimated from present-day individuals and therefore reflect present-day ancestry-specific allele frequencies. As these allele frequencies change through evolutionary time, it is possible to relate ancestries phylogenetically and make inferences about the common ancestors of ancestries. Projecting ancient individuals onto present-day ancestries will lead to increasingly incorrect inference as the age of the ancient individual increases. Thus, this issue is a bigger problem for Ice Age Europeans than for Bronze Age Europeans. This problem can be solved if allele frequencies for each of the ancestors of the present-day ancestries were known.

In summary, rather than three^3^ or four^8,10^ ancestral populations, we found considerably more population structure across 279 ancient Eurasians, involving a total of 18 autosomal ancestries, 12 Y DNA haplogroups, and 13 mitochondrial haplogroups, such that no sample was ancestrally homogeneous. Even if ancestries are inferred from extant individuals, ancestry analysis can provide historical insight in the absence of ancient DNA samples. Perhaps most importantly, using a consistent, unified nomenclature will enhance research of both ancient and present-day peoples.

## Methods

We retrieved and integrated data from 279 ancient Eurasians from 49 samples^3–5,8,10,12,18–21^. BAM files were processed using the program bam2mpg using a quality filter of 20. The global reference panel was previously described^7^. We performed supervised clustering by projecting the ancient Eurasians onto the 21 ancestries in this global reference panel using ADMIXTURE^2^. Standard errors were obtained from 200 bootstrap replicates. Inverse variance weighting was used to combine ancestry proportions across individuals within samples, accounting for both within- and between-individual variance. Assuming approximate normality, we induced sparsity by zeroing out any ancestry for which the 95% confidence interval included 0. Finally, the significant ancestry proportions were renormalized to sum to 1.

### Ethics

This project was determined to be excluded from IRB Review by the National Institutes of Health Office of Human Subjects Research Protections, Protocol 17-NHGRI-00282.

## Acknowledgements

This work utilized the computational resources of the NIH HPC Biowulf cluster (https://hpc.nih.gov). The contents of this publication are solely the responsibility of the authors and do not necessarily represent the official view of the National Institutes of Health. This research was supported by the Intramural Research Program of the Center for Research on Genomics and Global Health (CRGGH). The CRGGH is supported by the National Human Genome Research Institute, the National Institute of Diabetes and Digestive and Kidney Diseases, the Center for Information Technology, and the Office of the Director at the National Institutes of Health (Z01HG200362).

## Author Contributions

D.S. designed the project, performed the analysis, and wrote the manuscript.

